# Gaining Accuracy for Gene Expression Data by Parsimonious Models

**DOI:** 10.1101/2020.05.11.088484

**Authors:** Hugh G. Gauch, Yehao Zhang, Chang Chen

## Abstract

Gene expression data must be accurate in order to promote extensive, reliable, and repeatable results and to compare treatments with few false positives and false negatives. One way to gain accuracy is by advanced experimental techniques, and another is by good experimental design, including replication. But these may not be enough to achieve even one significant digit, as shown by an example using oat data. This article introduces an additional opportunity to increase accuracy that involves parsimonious models, which has not yet been implemented in the gene expression literature to the best of our knowledge. Basically, a parsimonious model gains accuracy by selectively recovering signal in its model while selectively relegating noise to a discarded residual. Typically, this accuracy gain is equivalent to averaging over several times as many replications, but its cost is trivial, merely some computation. Consequently, this neglected way to gain accuracy is quite cost effective. For gene expression research, accuracy gain by parsimonious models should be a standard component of best practices.

## Introduction

Gene expression data must be accurate in order to promote extensive, reliable, and repeatable results for vital applications in medicine, agriculture, and other areas of biology. Greater accuracy means that treatment comparisons have fewer false positives and false negatives. Accuracy has been gained through ongoing technological advances such as the recent 3’ RNA sequencing (Lohman et al. 2016), several adjustments applied to the original counts of transcripts (Li et al. 2017), and replication. Statistical analysis offers an additional way to gain accuracy that has been used to great advantage in countless applications in science and technology for over 50 years, but it has never yet been used in the gene expression literature to the best of our knowledge: parsimonious models. Implementing this additional opportunity as a standard component of best practices would be highly advantageous because current practices without parsimonious models produce rather inaccurate data.

The basic idea for accuracy gain by parsimonious models is to apply a model family with its sequence of increasingly complex (or decreasingly parsimonious) members to a dataset, and then to select the most predictively accurate member by cross validation. This best model is ordinarily relatively parsimonious, and it can be more accurate than the data used to construct it because it selectively recovers signal and discards noise (Gauch 2006, 2012:174–198, Fourdrinier et al. 2018).

The magnitude of the opportunity to gain accuracy by parsimonious models varies from application to application and from dataset to dataset. This opportunity increases with the size of a dataset since more information is helpful for fitting model parameters. It also increases with the noisiness of the data since exceedingly accurate data would offer little room for improvement. Precisely because gene expression datasets are typically large and noisy, they are ideal candidates for exceptionally large accuracy gains.

## Results

As a representative example of gene expression data, this article uses data on oats from Hu et al. (2019). The purpose of this experiment is to elucidate gene expression in developing hexaploid oat seeds so that oat breeders can release new cultivars with higher contents of health-promoting compounds, including avenanthramides that are reported to modulate signaling pathways associated with cancer, diabetes, inflammation, and cardiovascular disease.

Even after implementing advanced technology, applying numerous adjustments, and using replication, gene expression data can still have rather limited accuracy. For instance, the replicates for the first sample and first gene in the oat dataset are 7.587352171 and 5.627197681, which do not agree to even 1 significant digit. Presentation of these gene expression data with 10 digits may give an appearance of great accuracy, but discrepancy between these replicates on even their first digit quickly dispels that misperception. Indeed, of the 121 × 500 = 60500 replicate pairs in the oat dataset, only 37% agree to merely one significant digit, and only 10% to two significant digits. Likewise, for averages of 2 replicates, their root mean square (RMS) error is 17% of the RMS for the gene expression values, which again is indicative of less than one significant digit. Therefore, another opportunity to gain accuracy by parsimonious models merits serious consideration.

For gene expression data structured as a matrix, such as the oat example with its 121 samples by 500 genes, principal components analysis (PCA) provides a suitable model family consisting of PCA0, PCA1, PCA2, and so on with 0, 1, and 2 components included in the model before higher components are relegated to a discarded residual, until finally reaching the full model denoted by PCAF that retains all components, equals the data, and leaves no residual. Incidentally, PCA has been applied to gene expression data frequently, but for a different purpose, namely to produce low-dimensional graphs that display the main patterns in high-dimensional data (Lin et al. 2017, Brown et al. 2018).

A particular variant of PCA is ideal for gene expression studies because of special interest in *differential* gene expressions, especially between different treatments. Analysis of variance (ANOVA) applied to a 2-way factorial design of samples-by-genes recognizes three sources: sample main effects, gene main effects, and sample-by-gene (SxG) interaction effects. The SxG interactions just are these differential gene expressions, whereas both main effects are wholly irrelevant to differential responses of genes across samples. Therefore, the ideal variant is double-centered principal components analysis (DC-PCA), which is also called Additive Main effects and Multiplicative Interaction (AMMI; Gauch et al. 2008). It subtracts out both main effects before applying PCA to the SxG interactions, which alone carry information on differential responses of genes across samples. Because AMMI applies PCA to SxG exclusively, its principal components are distinguished by the name interaction principal components (IPCs). AMMI constitutes a model family, like all variants of PCA, comprised of AMMI0, AMMI1, AMMI2, and so on with 0, 1, and 2 IPCs included in the model before higher IPCs are relegated to a discarded residual, until finally reaching the full model denoted by AMMIF. The AMMI model includes all of the variation in the data: the sample main effects, gene main effects, and the SxG interaction effects. But AMMI uniquely analyzes these three sources of variation without confounding them (Gauch et al. 2008, Gauch et al. 2019). Because these three sources are of interest to researchers for different kinds of questions and reasons, AMMI is particularly appropriate.

Table 1 presents the ANOVA for AMMI analysis of the oat data. The degrees of freedom (df), sum of squares (SS), and mean square (MS) are listed for each source, and significance tests are done by ordinary F tests for treatment, sample, gene, and SxG effects and by FR tests for the IPCs (Piepho 1995). The sample main effects, gene main effects, and sample-by-gene (SxG) interaction effects account for 0.7%, 49.1%, and 50.2% of the treatment SS. The total SS for the SxG interactions for these oat data is spread across 120 IPCs. Table 1 shows that most of this variation is captured in just the first few IPCs. Indeed, IPC1 captures 68.3% of the interactions and is 8.7 times larger than the next IPC, so the differential gene expressions of special biological interest have an underlying structure that is remarkably simple and clear. The first 14 IPCs are statistically significant according to FR significance tests, and they account for 90.3% of the SxG interactions. This outcome can be explained by the amounts of signal and noise in the SxG interactions, and by the ability of AMMI to selectively capture signal in the early components that are retained and to selectively capture noise in the late components that are discarded. The SxG noise can be estimated by the error MS times the interaction df, 1.3606 59880 = 81472.7. By subtraction from the SxG total of 715840.7, the estimated SxG signal is 634368.0 or 88.6%, which is close to the 90.3% of the AMMI14 model.

**Table 1.** ANOVA for AMMI analysis of the oat data. The degrees of freedom (df), sum of squares (SS), and mean square (MS) are listed for each source, and significance tests are done by ordinary F tests for treatment, sample, gene, and SxG effects and by FR tests for the IPCs. The x total variation is partitioned into the treatment design and error; then the treatments are partitioned into sample main effects, gene main effects, and SxG interaction effects; and finally the SxG interactions are partitioned into 30 IPCs with the remaining IPCs 31 to 120 combined in x the residual. The first 14 IPCs are significant at the 0.001 level (***).

| Source | df | SS | MS | Probability |
| --- | --- | --- | --- | --- |
| Total | 120999 | 1509475.51 | 12.47511 |  |
| Treatment | 60499 | 1427159.05 | 23.58980 | 0.000000000000 *** |
| Sample | 120 | 10538.71 | 1404.36792 | 0.000000000000 *** |
| Gene | 499 | 700779.59 | 87.82257 | 0.000000000000 *** |
| S×G | 59880 | 715840.74 | 11.95459 | 0.000000000000 *** |
| IPC1 | 618 | 489053.67 | 791.34897 | 0.000000000000 *** |
| IPC2 | 616 | 55919.88 | 90.77902 | 0.000000000000 *** |
| IPC3 | 614 | 20054.59 | 32.66220 | 0.000000000000 *** |
| IPC4 | 612 | 12122.01 | 19.80720 | 0.000000000000 *** |
| IPC5 | 610 | 10556.14 | 17.30515 | 0.000000000000 *** |
| IPC6 | 608 | 10044.32 | 16.52026 | 0.000000000000 *** |
| IPC7 | 606 | 8159.17 | 13.46397 | 0.000000000000 *** |
| IPC8 | 604 | 7390.15 | 12.23534 | 0.000000000000 *** |
| IPC9 | 602 | 6806.98 | 11.30728 | 0.000000000000 *** |
| IPC10 | 600 | 6423.66 | 10.70611 | 0.000000000000 *** |
| IPC11 | 598 | 5655.98 | 9.45817 | 0.000000000000 *** |
| IPC12 | 596 | 5427.14 | 9.10594 | 0.000000000000 *** |
| IPC13 | 594 | 4839.04 | 8.14653 | 0.000000000000 *** |
| IPC14 | 592 | 4304.75 | 7.27153 | 0.000007657707 *** |
| IPC15 | 590 | 3937.95 | 6.69719 | 0.928799342886 |
| IPC16 | 588 | 3923.92 | 6.65071 | 0.999999999999 |
| IPC17 | 586 | 3582.52 | 6.11351 | 1.000000000000 |
| IPC18 | 584 | 3280.30 | 5.61694 | 1.000000000000 |
| IPC19 | 582 | 3184.41 | 5.47150 | 1.000000000000 |
| IPC20 | 580 | 2836.34 | 4.89025 | 1.000000000000 |
| IPC21 | 578 | 2546.57 | 4.42112 | 1.000000000000 |
| IPC22 | 576 | 2526.97 | 4.37192 | 1.000000000000 |
| IPC23 | 574 | 2409.78 | 4.19823 | 1.000000000000 |
| IPC24 | 572 | 2257.19 | 3.94614 | 1.000000000000 |
| IPC25 | 570 | 1951.27 | 3.42329 | 1.000000000000 |
| IPC26 | 568 | 1750.19 | 3.08131 | 1.000000000000 |
| IPC27 | 566 | 1743.41 | 3.08024 | 1.000000000000 |
| IPC28 | 564 | 1596.73 | 2.83109 | 1.000000000000 |
| IPC29 | 562 | 1427.25 | 2.53960 | 1.000000000000 |
| IPC30 | 560 | 1240.34 | 2.21489 | 1.000000000000 |
| Residual | 42210 | 28888.12 | 0.68439 | 1.000000000000 |
| Error | 60500 | 82316.46 | 1.36060 |  |

Table 2 shows the results from cross validation, including the statistical efficiency of the AMMI model family, with AMMI14 being the most predictively accurate model and achieving a statistical efficiency of 3.20. There are 2 replications, so cross validation selects one replicate at random for the model data and the other for the validation data, using a separate randomization for each of the 60500 matrix entries. This process was repeated 50 times and the results were averaged for greater accuracy. The initial result from cross validation is the prediction variance, namely the mean squared discrepancy between model predictions and validation observations. The prediction variance for AMMI14 is 1.78633. For comparison, the prediction variance for AMMIF or the actual data is the error mean square 1.36060 for the single observations in the model data plus the same for the validation data, namely 2.72120. Since 1.78633 is decidedly smaller than 2.72120, cross validation provides empirical proof that the AMMI14 model is more accurate than the data used to construct this model. The AMMI14 predictions and the validation observations are discrepant because both are imperfect. The model variance can be obtained by subtraction, 1.78633 – 1.36060 = 0.42573. Finally, by definition, the statistical efficiency equals the variance of the full model divided by the variance of a given model, namely 1.36060 / 0.42573 = 3.20 for the AMMI14 model. For comparison, the statistical efficiency of the full model AMMIF, which equals the averages over replicates, is automatically equal to 1. Note that the diagnosis of AMMI14 by cross validation agrees with the diagnosis from FR significance tests given earlier, and again this result can be explained by the amounts of signal and noise in the SxG interactions, so this diagnosis is solid.

**Table 2.** Predictive accuracy and statistical efficiency of the AMMI model family. Results are given for AMMI0 to AMMI30 and the full model AMMIF, which is AMMI120, applied to oat gene expression data. The prediction variance is the mean squared discrepancy between model predictions and validation observations, obtained by cross validation. The variance of the validation observations is simply the error mean square. The variance of a model equals its prediction variance minus its validation variance. The statistical efficiency equals the model variance of the full model divided by that for the given model. The most predictively accurate model is AMMI14, marked with an asterisk. Its statistical efficiency of 3.20 means that this model is as accurate as would be averages based on 3.20 times as many replications.

| AMMI Model | Prediction Variance | Validation Variance | Model Variance | Statistical Efficiency |
| --- | --- | --- | --- | --- |
| AMMI0 | 6.62024 | 1.36060 | 5.25964 | 0.26 |
| AMMI1 | 2.59276 | 1.36060 | 1.23216 | 1.10 |
| AMMI2 | 2.15062 | 1.36060 | 0.79002 | 1.72 |
| AMMI3 | 2.01070 | 1.36060 | 0.65010 | 2.09 |
| AMMI4 | 1.93492 | 1.36060 | 0.57432 | 2.37 |
| AMMI5 | 1.88133 | 1.36060 | 0.52073 | 2.61 |
| AMMI6 | 1.82623 | 1.36060 | 0.46563 | 2.92 |
| AMMI7 | 1.80058 | 1.36060 | 0.43998 | 3.09 |
| AMMI8 | 1.80616 | 1.36060 | 0.44556 | 3.05 |
| AMMI9 | 1.81902 | 1.36060 | 0.45842 | 2.97 |
| AMMI10 | 1.80275 | 1.36060 | 0.44215 | 3.08 |
| AMMI11 | 1.79985 | 1.36060 | 0.43925 | 3.10 |
| AMMI12 | 1.78810 | 1.36060 | 0.42750 | 3.18 |
| AMMI13 | 1.78774 | 1.36060 | 0.42714 | 3.19 |
| AMMI14 | 1.78633 | 1.36060 | 0.42573 | 3.20 * |
| AMMI15 | 1.78954 | 1.36060 | 0.42894 | 3.17 |
| AMMI16 | 1.79852 | 1.36060 | 0.43792 | 3.11 |
| AMMI17 | 1.80321 | 1.36060 | 0.44261 | 3.07 |
| AMMI18 | 1.80420 | 1.36060 | 0.44360 | 3.07 |
| AMMI19 | 1.80337 | 1.36060 | 0.44277 | 3.07 |
| AMMI20 | 1.80338 | 1.36060 | 0.44278 | 3.07 |
| AMMI21 | 1.80558 | 1.36060 | 0.44498 | 3.06 |
| AMMI22 | 1.81038 | 1.36060 | 0.44978 | 3.03 |
| AMMI23 | 1.81694 | 1.36060 | 0.45634 | 2.98 |
| AMMI24 | 1.82604 | 1.36060 | 0.46544 | 2.92 |
| AMMI25 | 1.83430 | 1.36060 | 0.47370 | 2.87 |
| AMMI26 | 1.84251 | 1.36060 | 0.48191 | 2.82 |
| AMMI27 | 1.85240 | 1.36060 | 0.49180 | 2.77 |
| AMMI28 | 1.86366 | 1.36060 | 0.50306 | 2.70 |
| AMMI29 | 1.87772 | 1.36060 | 0.51712 | 2.63 |
| AMMI30 | 1.89247 | 1.36060 | 0.53187 | 2.56 |
| AMMIF | 2.72120 | 1.36060 | 1.36060 | 1.00 |

Figure 1 shows the relationship between model complexity and predictive accuracy—a response called Ockham’s hill (MacKay 1992, Gauch 2006). The AMMI0 to AMMI30 models along the abscissa increase in complexity to the right, or increase in parsimony or simplicity to the left. The ordinate shows statistical efficiency, which is a measure of predictive accuracy. To the left of the peak, excessively simple models underfit real signal; to the right, excessively complex models overfit spurious noise. At one extreme, AMMI0 captures none of the SxG interactions, which means none of the signal (which is bad) and none of the noise (which is good); whereas at the opposite extreme, AMMIF captures all of the SxG interactions, which means all of the signal (which is good) and all of the noise (which is bad). Ockham’s hill shows that it is better to have fairly small problems with both underfiting signal and overfiting noise than it is to have a huge problem with either underfiting or overfiting. Not shown are higher members of this family from AMMI31 to AMMI120, which is the full model AMMIF. Their statistical efficiencies decline to an asymptote of 1 for the full model. Incidentally, although AMMI14 is most accurate, the peak on this Ockham’s hill happens to be rather broad, and all models from AMMI7 through AMMI22 achieve a statistical efficiency over 3, so all of them are nearly optimal and all are much more accurate than the data used to construct these models.

**Fig 1.**
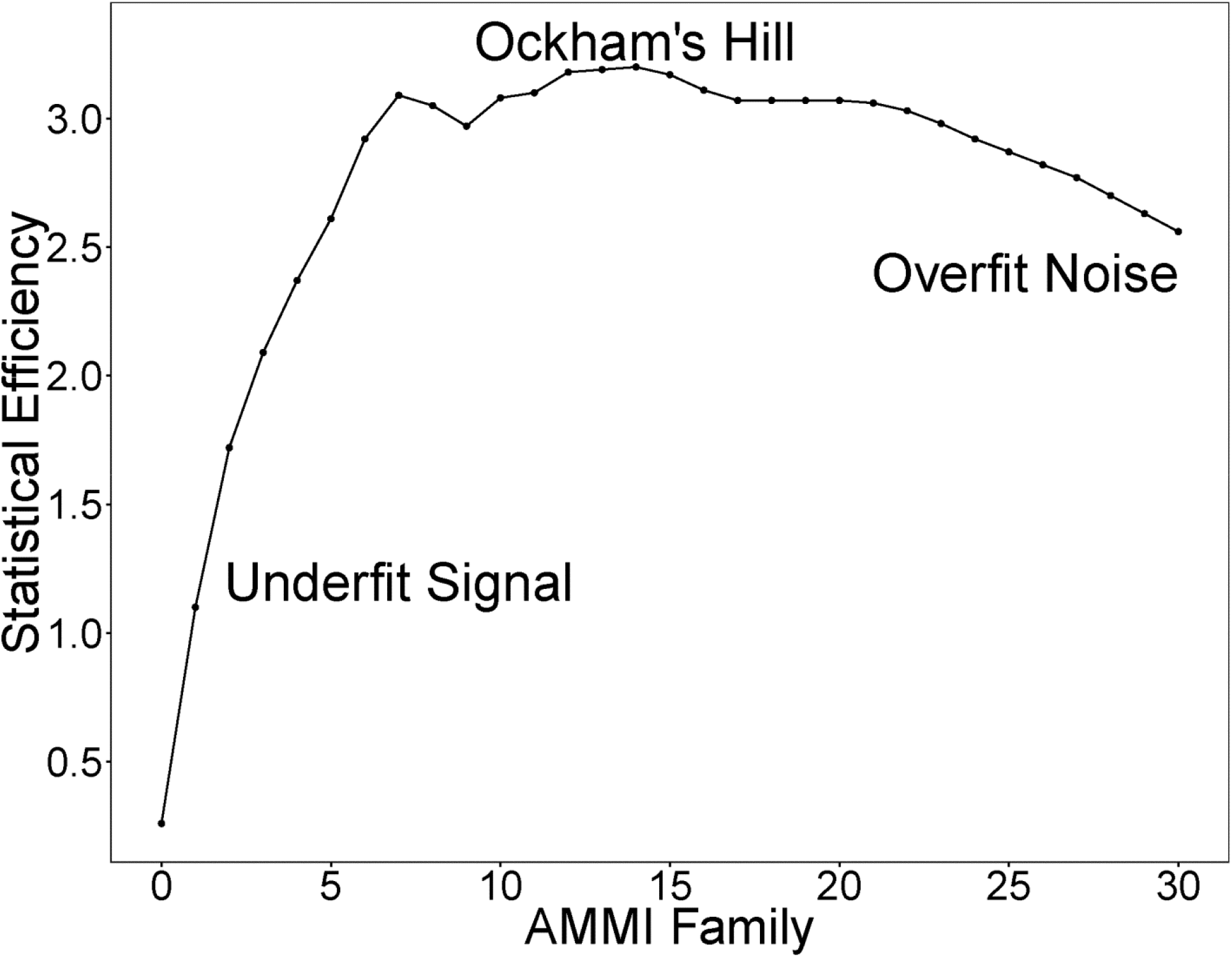
Ockham’s hill for the oat gene expression data. The AMMI model family is shown from AMMI0 to AMMI30, which includes the peak of Ockham’s hill at a statistical efficiency of 3.20 for AMMI14. To the left of the peak, models are less accurate because they underfit signal, and to the right because they overfit noise.

## Discussion

Replicated data are ideal for gaining accuracy by parsimonious models because statistical efficiency can be quantified and Ockham’s hill displayed, but unreplicated data can still function to a limited and yet useful degree. AMMI analysis does not require replication, so the SSs or eigenvalues of the IPCs are still available. A rough estimate of the error mean square from similar but replicated experiments is all of the additional information that is needed to estimate the amounts of signal and noise in the SxG interactions, and thereby to diagnose the most predictively accurate member of the AMMI model family. That said, replication is a hallmark of scientific research for many good reasons, and fortunately replication is a reasonably common feature of gene expression experiments.

An important distinction is that some experimental designs adjust estimates closer to their true values, but others do not. For example, randomized complete block designs can reduce pure error and thereby increase statistical significance, yet they do not adjust estimates. By contrast, randomized incomplete block designs do both, they reduce pure error *and* adjust estimates closer to their true values. Whenever both the experimental and treatment designs offer opportunities to gain accuracy, adjust estimates with the experimental design first, and then supply those improved values for AMMI analysis.

The customary procedure has been to diagnose the most predictively accurate member of the AMMI model family using cross validation (or F_R_ tests or signal-and-noise considerations), and then to apply that model to the recombined data using all replicates. But an open question remains about how to estimate the statistical efficiency of that final result. The usual procedure applied to the oat data would be to multiply the statistical efficiency of 3.20 from modeling 1 replicate by the number of replications in the final model, 3.20 × 2 = 6.40. However, this gives an overestimate of the statistical efficiency because averages over 2 replicates are more accurate than a single replicate, and more accurate data present parsimonious models with a smaller opportunity to gain accuracy. Furthermore, more accurate data can support more AMMI parameters, so there may be a shift to the most accurate model being a somewhat higher member of the AMMI model family—although that shift is rather inconsequential in the common case that Ockham’s hill is rather broad. Meanwhile, until this open question has been answered, it seems more important to take full advantage of all of the data in the final model than to have cross validation available for quantifying the statistical efficiency.

It is axiomatic that data having greater accuracy support more extensive and reliable results. Indeed, this is precisely why the usual normalizations and adjustments that are applied to the original transcript counts or other raw data in gene expression research have been standard components of best practices, and also is why accuracy gain by parsimonious models deserves to become another component of best practices. However, the extent of additional biological insight and research progress that would emerge from parsimonious models is an open question that requires empirical investigation with several representative datasets. It would be interesting to compare biological conclusions based on three versions of gene expression data: the original transcript counts or raw data, the estimates after the usual adjustments, and the estimates after both the usual adjustments and parsimonious models. Although this brief article does not include that extensive empirical investigation, it does provide methodology and motivation to address this open question about biological dividends.

## Conclusion

Only a single example using oat data was presented in this article, but a simple argument shows that the opportunity to gain accuracy by parsimonious models applies to gene expression research in great generality. For starters, cross validation provides empirical proof for this oat example that a parsimonious model can gain accuracy. Furthermore, statistical theory provides an explanation of accuracy gain in terms of selective recovery of signal in early model parameters as described by Ockham’s hill, and it shows why this selectivity works especially well for large and noisy datasets. These two dataset properties constitute the necessary and sufficient conditions for substantial accuracy gain, and typically gene expression datasets are large and noisy. Therefore, for gene expression research, accuracy gain by parsimonious models should be a standard component of best practices.

## Methods

### Data

Hu et al. (2019) describes their methods for gene expression data on oats in detail, but basically they used 3’ mRNA sequencing, counted the number of reads mapped onto a gene, and then made several adjustments. Their expression matrix of 59815 transcripts by 397 samples was included as Appendix S2 in their supporting information. Their Figure 2 used a subset with the 500 transcripts with highest variance and the 326 samples with more than 0.5 million mapped reads, and they kindly shared that dataset with us. Finally, we produced a subset with no missing replications that has 121 samples, 500 genes, and 2 replications (where we are using the word “samples” to mean individual treatments, and each of these samples has 2 replicates).

### Software

ANOVA, PCA, and cross validation are all common statistical analyses that are readily available in numerous statistical packages. We found it convenient, however, to write our own software, and include our R code in the supporting information.

## Supporting information

**S1 Table. The oat gene expression data as received from Haixiao Hu with the 500 transcripts with highest variance and the 326 samples with more than 0.5 million mapped reads.** Haixiao Hu granted permission for us to include these data as supporting information in this article.

(XLSX)

**S1 Text. Our subset of the received oat gene expression data with no missing replications that has 121 samples, 500 genes, and 2 replications, formatted as required for our software.**

(TXT)

**S2 Text. The AMMI14 estimates for the matrix of 121 samples by 500 genes, formatted as required for our software.**

(TXT)

**S3 Text. The R code for AMMI analysis, ANOVA tables, and cross validation.** This R code was produced for our own in-house research purposes, not as polished and public software, but it is made available here for the sake of transparency in research.

(TXT)

## Acknowledgements

Haixiao Hu kindly made available to us the gene expression data that was used for Figure 2 in the cited 2019 article in *Plant Biotechnology Journal*, and he also granted permission for us to include these data in our supporting information. We appreciate helpful suggestions on earlier drafts of this manuscript from Haixiao Hu, Hans-Peter Piepho, Sheng Qian, and Mark Sorrells.

## Author contributions

**Conceptualization:** Hugh G. Gauch, Jr.

**Data curation:** Chang Cheng and Yehao Zhang

**Software:** Chang Cheng

**Visualization:** Yehao Zhang and Chang Cheng

**Supervision:** Hugh G. Gauch, Jr.

**Writing – original draft:** Hugh G. Gauch, Jr.

**Writing – review & editing:** Hugh G. Gauch, Jr., Yehao Zhang, and Chang Cheng

## Notes

### Competing Interest Statement

The authors have declared no competing interest.

